# Reconstruction of clone- and haplotype-specific cancer genome karyotypes from bulk tumor samples

**DOI:** 10.1101/560839

**Authors:** Sergey Aganezov, Benjamin J. Raphael

**Affiliations:** Department of Computer Science, Princeton University, Princeton, NJ 08540; Department of Computer Science, Johns Hopkins University, Baltimore, MD 21218

## Abstract

Many cancer genomes are extensively rearranged with highly aberrant chromosomal karyotypes. These genome rearrangements, or structural variants, can be detected in tumor DNA sequencing data by abnormal mapping of sequence reads to the reference genome. However, nearly all cancer sequencing to date is of bulk tumor samples which consist of a heterogeneous mixture of normal cells and subpopulations of cancers cells, or clones, that harbor distinct somatic structural variants. We introduce a novel algorithm, <u>R</u>econstructing <u>C</u>ancer <u>K</u>aryotypes (`RCK`), to reconstruct haplotype-specific karyotypes of one or more rearranged cancer genomes, or clones, that best explain the read alignments from a bulk tumor sample. `RCK` leverages specific evolutionary constraints on the somatic mutation process in cancer to reduce ambiguity in the deconvolution of admixed DNA sequence data into multiple haplotype-specific cancer karyotypes. In particular, `RCK` relies on generalizations of the infinite sites assumption that a genome re-arrangement is highly unlikely to occur at the same nucleotide position more than once during somatic evolution. `RCK`’s comprehensive model allows us to incorporate information both from short and long-read sequencing technologies and is applicable to bulk tumor samples containing a mixture of an arbitrary number of derived genomes. We compared `RCK` to the state-of-the-art method `ReMixT` on a dataset of 17 primary and metastatic prostate cancer samples. We demonstrate that `ReMixT`’s limited support for heterogeneity and lack of evolutionary constrains leads to reconstruction of implausible karyotypes. In contrast, `RCK`’s infers cancer karyotypes that better explain read alignments from bulk tumor samples and are consistent with a reasonable evolutionary model. `RCK`’s reconstructions of clone- and haplotype-specific karyotypes will aid further studies of the role of intra-tumor heterogeneity in cancer development and response to treatment. `RCK` is available at https://github.com/raphael-group/RCK.

## 1 Introduction

The somatic mutations that drive cancer development range across all genomic scales, from single nucleotide mutations through large-scale genome rearrangements [54, 20, 58, 47]. Whole-genome sequencing of tumor samples has enabled the detection of all classes of somatic mutations; however, specialized algorithms are required to identify each class of mutations from the short DNA sequence reads obtained by current technologies [31, 37, 28, 49, 30, 60]. In addition, nearly all cancer sequencing to date has been of bulk tumor tissue, which is generally a mixture of normal (noncancerous) cells and (sub)populations of cancerous cells, or *clones*, that often are not genetically identical. Quantifying this *intra-tumor heterogeneity* is essential for understanding the processes that drive cancer development and also helps inform treatment strategies [2, 44, 34].

Here we consider the problem of describing the large-scale organization of one or more cancer genomes that are derived form a normal human reference genome via large-scale rearrangements. The large-scale organization of a cancer genome is described by two features. First, is the number of copies of each segment of the genome. Many methods (e.g. [57, 9, 7, 40, 24, 19, 43, 63]) have been developed to identify copy number values for heterogeneous, bulk tumor samples. Second, are genome rearrangements (e.g. chromosomal inversions and translocations) that link together distant segments of the normal genome. Many methods have been developed to predict such *novel adjacencies* (e.g. [51, 48, 30, 10, 60, 49, 16, 52, 66, 50, 27, 17]. However, these methods do not distinguish between adjacencies from different homologous chromosomes or from different cancer clones within a bulk sample; i.e. they assume the human genome is *haploid reference* and that the tumor is homogeneous.

A more challenging problem is to combine and reconcile the information about segment copy numbers and novel adjacencies into genome *karyotypes*, or the alignment of cancer genome and the healthy genome that depicts the number of occurrences of every segment in the cancer genome, and the adjacencies between these segments on the cancer genome. Multiple methods have been developed to solve some variations of this cancer genomes karyotype reconstruction problem including [42, 32, 46, 35, 14, 11, 15]. However, each of these methods rely on simplifying assumptions that do not adequately address the challenges in real cancer sequencing data. For example, `SVclone` [11] focuses solely on inferring genome-specific copy numbers for novel adjacencies, without attempting to reconstruct complete karyotypes of the derived genomes. `PREGO` [42] and `Karyotype Reconstruction` [15] assume that the human reference genome is haploid, thus losing important information about alleles involved in rearrangements. `Weaver` [32, 46] assumes that the cancer sample contains only a single derived genome (with a possible admixture of the reference genome), and lacks a proper support of reciprocal novel adjacencies, which can emerge both from copy number neutral somatic rearrangements (e.g., inversions, balanced translocations, etc), as well as from more complex “catastrophic rearrangements” such as chromoplexy and chromothripsis [53, 6, 26, 4, 61, 41]. `ReMixT` [35] allows for tumor heterogeneity, but fixes the number of derived genomes in the observed cancer sample to 2. Moreover, while `ReMixT` aims to infer genome- and allele-specific segment copy numbers for a 2-genome sample (with a possible admixture of the reference genome), the genome-specific copy numbers for novel adjacencies that are inferred by `ReMixT` lack information about which homologous copies of the segments are actually involved in observed novel adjacencies. Lastly, `Weaver`, and `ReMixT` produce karyotypes with biologically unlikely scenarios where rearrangements occur repeatedly at the same homologous loci in different cancer clones. We summarize these limitations of existing methods in Table S1.

Here we propose a novel algorithm, <u>R</u>econstructing <u>C</u>ancer <u>K</u>aryotypes (`RCK`), for deriving the karyotypes of cancer genomes in a heterogeneous tumor sample from next-generation (and 3rd-generation, when available) sequencing data. `RCK` distinguishes itself from existing methods by several features including: (i) support for diploid reference genome distinguishing between alleles of segment copy numbers and novel adjacencies (ii) joint inference of both segment and adjacency copy numbers in both clone- and haplotype-specific fashion; (iii) comprehensive support for sample heterogeneity ranging from homogeneous samples with a single derived genome to heterogeneous samples with an arbitrary number of clones; (iv) enforcement of somatic evolutionary constraints on all genomes within a sample; (v) unique ability to incorporate groups of novel adjacencies from 3rd-generation sequencing technologies into the inference model. We demonstrate the advantages of `RCK` by comparing its performance to `ReMixT` on a dataset of 17 primary and metastatic prostate cancer samples. We find that `RCK` infers more plausible karyotypes that conform to an evolutionary model and have allele-specific segment copy numbers that agree with leading copy number inference algorithms.

## 2 Results

### 2.1 RCK algorithm

We introduce <u>R</u>econstructing <u>C</u>ancer <u>K</u>aryotypes (`RCK`), an algorithm to construct the large-scale organization of one or more cancer genomes present in a bulk tumor sample. Each cancer genome in the sample arises from a sequence of somatic genome rearrangements and copy number number aberrations that transform a healthy normal genome into a cancer genome. As a result of these somatic mutations, each cancer genome can be represented as a *karyotype graph* – or more briefly a *karyotype*. A karyotype graph includes: (1) a collection of contiguous *segments* from the human reference genome, each segment with a label (`A` or `B`) distinguishing the two homologous chromosomes; an integer *copy number* for each segment; (3) a collection of *adjacencies* that join the ends of segments; (4) an integer copy number for each adjacency. The karyotype graph describes an alignment between the cancer genome and healthy genome (analogous to the breakpoint graph [1, 3] in genome rearrangement studies). The karyotype graph also represents the information about the cancer genome sequence that can be inferred from DNA sequencing technologies whose reads lengths are shorter than the length of genome rearrangements.

`RCK` solves the following *Cancer Karyotype Reconstruction Problem*: given allele-specific segment copy numbers and a list of *novel adjacencies* (i.e. pairs of genomic loci that are measured as adjacent in the cancer genome, but distant in the normal reference) from a bulk tumor sample, derive karyotype graph(s) for the cancer genome(s) present in the tumor sample. Several challenges emerge in the development of an algorithm to solve this problem. The first challenge is that the many methods for inferring allele-specific copy numbers from bulk tumor sequencing data (e.g. [57, 9, 7, 40, 24, 19, 35, 63]) do not preserve the allelic information across multiple adjacent segments. Specifically, these methods output a pair of copy number vectors, ĉ = [*ĉ*_1_, *ĉ*_2_, …,*ĉ*_*m*_] and č = [*č*_1_, *č*_2_, …, *č*_*m*_], where the pair (*ĉ*_*j*_, *č*_*j*_) ∈ ℕ^2^ indicates the number of copies of each of the two homologous copies of segment *j* from the reference genome that are present in the cancer genome. However, each of these pairs are *unordered*: it is not known whether *ĉ*_*j*_ is the number of segment from the maternal chromosome or the paternal chromosome; moreover, the identification of *ĉ*_*j*_ as maternal or paternal is independent for each *j*.

The second challenge results is that the many methods for inferring *novel adjacencies* from bulk tumor sequencing data [51, 48, 30, 10, 60, 49, 16, 52, 66, 50, 27, 17] generally do not include two important attributes in their output: (i) the alleles (maternal or paternal) that are joined by the adjacency; (ii) the copy number(s) of the adjacency in each genome in the sample. Because of this incomplete information in the allele-specific copy numbers and novel adjacencies, cancer genome karyotypes are not directly available.

`RCK` derives optimal cancer genome karyotype(s) from allele-specific copy numbers and novel adjacencies by solving an optimization problem on a graph, called the *Diploid Interval Adjacency Graph* (DIAG) (Figure 1). The vertices of the DIAG are *extremities*, or the positions in the human reference genome of the endpoints of the segments that are rearranged to form the cancer genomes present in the sample. Specifically, we enumerate the segments of the reference genome 1, …, *m*. Each segment *j* has the form 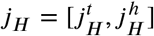, where 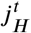 and 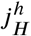 are extremities. The label *t* indicates that the extremity is the *tail*, or starting coordinate of the segment in the reference genome, while the label *h* indicates the head, or ending coordinate in the reference genome. A *haplotype* label *H* ∈ {`A`, `B`} indicates which copy of the two homologous chromosomes in the reference (`A` or `B`) is the source of the segment. Adjacent extremities of consecutive segments that follow each other along the chromosome in the genome constitute an *adjacency*. We distinguish between two types of adjacencies: *reference adjacencies* that are present in the reference genome, and *novel adjacencies* that are *not* present in the reference genome. Thus, the DIAG has three types of edges: (1) *segment edges* 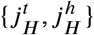 join extremities from a segment; (2) *reference adjacency edges* 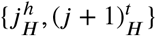 join extremities of adjacent segments on the reference genome; (3) *novel adjacency edges* 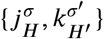, where *H, H′* ∈{`A,B`}, and, *σ, σ ′* ∈ {*t, h*}. Importantly, since a measured novel adjacency 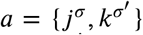 does not generally include allelic information, we add all 4 possible labeled versions of the adjacency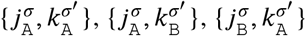, and 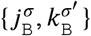 to the DIAG.

**Figure 1:**
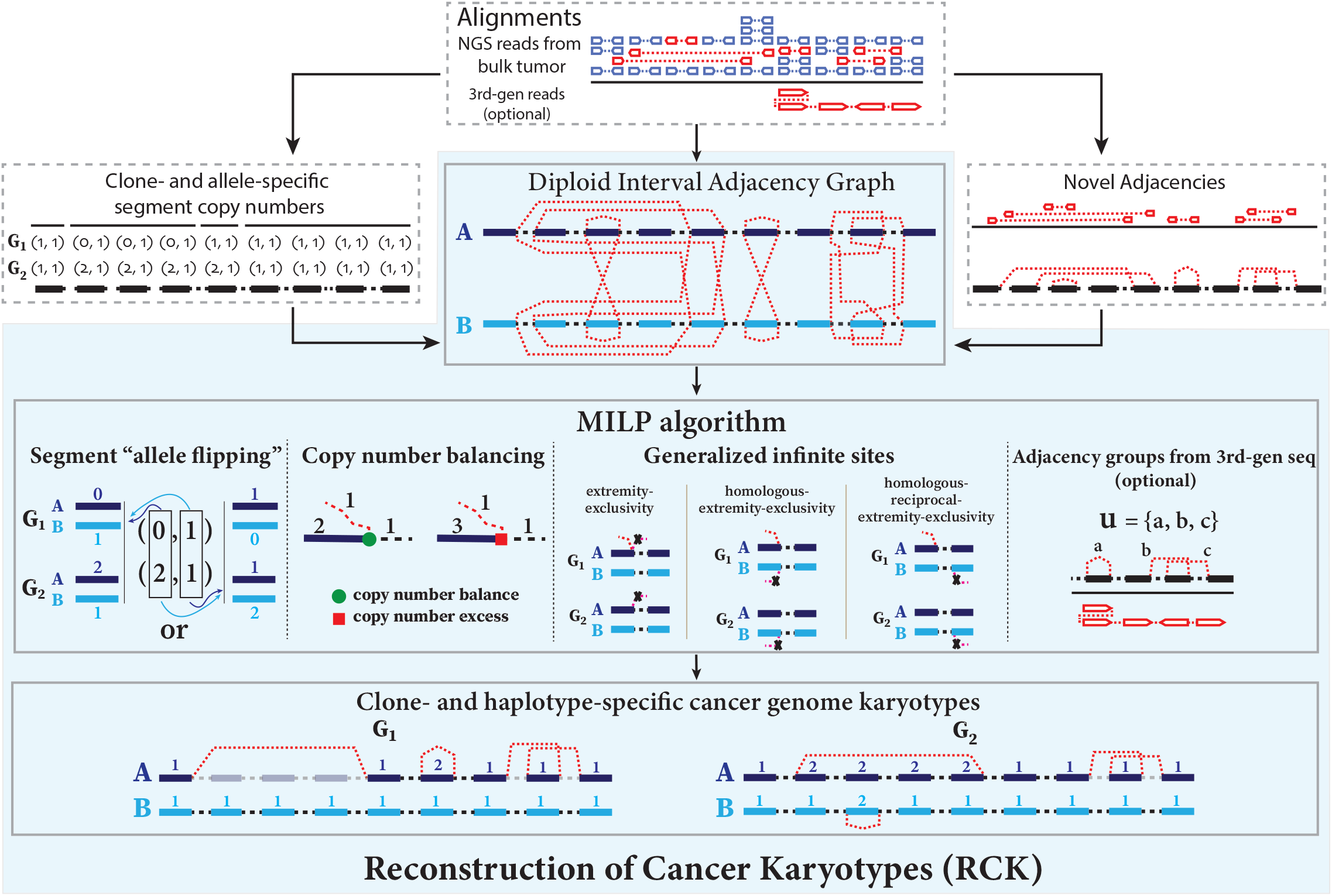
Overview of the RCK algorithm. Read alignments from bulk tumor sample are input to existing algorithms to identify clone- and allele-specific segment copy numbers (left) and novel adjacencies (right). The `RCK` algorithm (blue shaded elements) builds a *diploid interval adjacency graph* integrating copy number and novel adjacency infor an mixed-integer linear program (MILP) that finds an optimal assignment of copy numbers and novel adjacencies to alleles and clones, subject to copy number balance on segment ends and satisfying evolutionary constraints from a generalized infinite sites model. Constraints on groups of novel adjacencies from the 3rd generation sequencing technologies may optionally be included. The output of `RCK` are clone- and haplotype-specific cancer genome karyotypes.

A chromosome in the cancer genome corresponds to a walk in the DIAG that alternates between segment edges and reference/novel adjacency edges, and where the number of times every segment/adjacency edge is visited encodes the respective segment/adjacency copy number (see Methods 4.2). Thus, all vertices (except telomere vertices) should satisfy the *copy number balance condition*: the copy number of the incident segment edge equals the sum of the copy numbers of the incident reference edge and novel adjacency edge(s).

The Cancer Karyotype Reconstruction Problem thus can be formulated as the problem of finding an edge multiplicity *µ*_*G*_(*e*) for each edge *e* and each cancer genome *G* such that: (i) each extremity (vertex *v*) satisfies the *copy number balancing* conditions (Equations (9), (10) in Methods); (ii) the copy numbers *µ*_*G*_(*j*_*A*_) and *µ*_*G*_(*j*_*B*_) of homologous segments *j*_*A*_ and *j*_*B*_ are approximately equal to the allele-specific copy numbers (*ĉ*_*j*_ and *č*_*j*_); (iii) most of the novel adjacencies are present in at least one genome (i.e. *µ*_*G*_ (*e*) ≥ 0 for novel adjacency edge *e* in at least one genome *G*).

A major difficulty with the above formulation of the Cancer Genome Karyotype Reconstruction Problem is that there are often numerous solutions, many of which are biologically implausible. Considerable ambiguity arises from the lack of `A` /`B` labels on the measured novel adjacencies. The lack of allelic label means that each measured novel adjacency corresponds to 4 edges in the DIAG. However, selecting one of these four possible *allele-specific* novel adjacencies *independently* for each measured novel adjacency is unwise. Rather, the somatic evolutionary process imposes several constraints on the possible structures of inferred karyotypes. In particular, we derive conditions on allowed novel adjacencies from the *infinite sites (IS) assumption* commonly used in evolutionary studies. The infinite sites assumption is that a mutation does not occur at the same *locus* more than once during the course of evolution. The locus of a single-nucleotide mutation is readily defined as a genomic position. However, the locus for a large-scale genome rearrangement is not apparent, and could be defined as either (or both) of the genomic positions of the extremities in the adjacency as well as adjacent genomic positions of “reciprocal” extremities. We define multiple constraints on the extremities that may be involved in novel adjacencies (Figure 1). These constraints generalize the infinite sites assumption to the case of multiple genomes that are derived from a diploid reference genome by a sequence of large-scale genome rearrangements. First, **extremity-exclusivity** is the constraint that an extremity is involved in *at most one* novel adjacency. Second, **homologous-extremity-exclusivity** is the constraint that an extremity and its homolog *cannot both* be involved in a novel adjacency. Third, **homologous-reciprocal-extremity-exclusivity** is the constraint that an extremity and its reciprocal mate of the homologous chromosome *cannot both* be involved in a novel adjacency. All of these constraints are natural generalizations of the infinite sites assumption; however, they have not been distinguished consistently in previous publications (See Methods). As a result, previous methods can yield implausible genome reconstructions, as we will demonstrate below.

`RCK` solves the optimization problem of finding edge multiplicities *µ*(*e*) satisfying conditions (i), (ii), and (iii) above and *also* where the novel adjacencies inferred to be present (*µ*_*G*_(*e*) *>* 0) satisfy the generalized infinite sites constraints jointly across all clones. We solve this problem using a mixed-integer linear program (see Supplement S2.2). `RCK` also allows for grouping of novel adjacencies that are measured to be present on the same cell or longer read when such information is available from 3rd generation sequencing technologies (e.g. single cell sequencing, linked read sequencing [66, 52, 16], or long read sequencing [49, 17, 50, 27]). See Methods 4.4 for further details.

### 2.2 Evaluation and comparison of RCK

We compare `RCK` to `ReMixT`, the only other existing method which both derives multiple tumor clones from bulk sequencing data and distinguishes between homologous chromosomes. `ReMixT` takes read alignments and novel adjacencies as input and infers clone- and allele-specific copy numbers for segments, as well as clone-specific copy numbers for novel adjacencies. Importantly, `ReMixT` does *not* infer haplotype `A` /`B` labels for the extremities that are involved in each novel adjacency. We will show below that this lack of assignment of each novel adjacency to a homologous chromosome leads to unusual genome reconstructions in many cases.

#### 2.2.1 Data processing

We analyze a cancer sequencing dataset from Gundem et al. [23], which consists of whole-genome sequencing data from 49 samples from 10 metastatic prostate cancer patients. Segment copy numbers inferred by `Battenberg` [40] were obtained from the publication [23] and read alignments for every sample were obtained from the authors. For each sample, `Battenberg` output includes: (i) the number of clones; (ii) allele-specific copy numbers for each genomic segment in each clone; (iii) the occurrence of a whole genome duplication (WGD) when reported tumor ploidy *>* 3. We also used `HATCHet` [63], a recently developed algorithm that infers allele-specific copy numbers for one or more cancer clones as well as the presence of WGD by joint analysis of multiple sequenced samples from the same patient. We considered the 17 samples where both `Battenberg` and `HATCHet` agreed on the number of clones present.

For novel adjacencies, we used the predictions from `brass2` (https://github.com/cancerit/BRASS), which we obtained from the the original publication [23]. `brass2`, like most methods that identify novel adjacencies from aligned DNA sequence reads, has some uncertainty in the exact genomic coordinate involved in a novel adjacency. This uncertainty can be an issue when determining whether an adjacency is part of a reciprocal event (e.g. inversion or reciprocal translocation). Thus, we adjusted the coordinates of extremities to obtain refined coordinates for loci involved in reciprocal novel adjacencies. For `RCK`, we also aligned the positions of extremities of segments from `Battenberg` or `HATCHet` to the positions of extremities from novel adjacencies determined by `brass2`. See Methods 4.6 for further details.

We divided the cancer samples into two groups according to the number of tumor clones predicted by both `Battenberg` and `HATCHet` : *homogeneous* samples containing only one tumor clone (samples `A21g`, `A21h`, `A24c`, `A24d`, `A24e`, `A34a`, `A34c`, `A34d`); and *heterogeneous* samples containing two tumor clones (samples `A10c`, `A12c`, `A12d`, `A17d`, `A31a`, `A31d`, `A31e`, `A31f`, `A32e`). Notably, there was only one sample (`A12c`) where `Battenberg` and `HATCHet` disagreed on the presence of a WGD.

For each sample, we ran `RCK` requiring that: (1) the only telomeres in the inferred cancer genomes are telomeres from the reference genome (i.e. extremities that are not the endpoints of reference chromosomes have copy number balance); (2) at least a fraction `P` of the input novel adjacencies are present in at least one of the derived genomes in a sample, for `P` = 1.0, 0.9, 0.75, 0.5. `ReMixT` does not allow control over telomeres or the fraction of novel adjacencies, and thus we ran `ReMixT` using default parameters.

#### 2.2.2 Heterogeneous tumor samples

We first compared the allele-specific segment copy numbers inferred by `ReMixT`and the haplotype-specific segment copy numbers inferred by `RCK`to the allele-specific copy numbers from `HATCHet`and `Battenberg`, using a length-weighted segment copy number distance (equation (15) in Methods). We found that in all but two cases (samples `A10c`and `A12c`with a NAs utilization parameter `P`= 1.0), the segment copy numbers inferred by `RCK` are more similar to the copy numbers from `HATCHet`(Figure 2A) and `Battenberg`(Figure S2A). We also observed that `RCK`’s ability to control the fraction of input novel adjacencies that are required to be utilized in the inferred karyotype (Methods 4.3, 4.5) allows for more plausible reconstructions: the distance between input copy numbers is largest when we require `RCK`to use all novel adjacencies, but the distance decreases and stabilizes when a small fraction of novel adjacencies are excluded (`P`0.9). We note that the largest distances between `ReMixT`and `HATCHet`(or `Battenberg`) inferred copy numbers are on four samples (`A31a`, `A31d`, `A31e`, and `A31f`) where both `Battenberg`and `HATCHet`inferred a WGD. In these four samples, the high segment copy number values output by `ReMixT`also suggest many copy number changes; however, the large distances indicate that these inferred copy numbers may not align well with copy numbers expected from a WGD.

**Figure 2:**
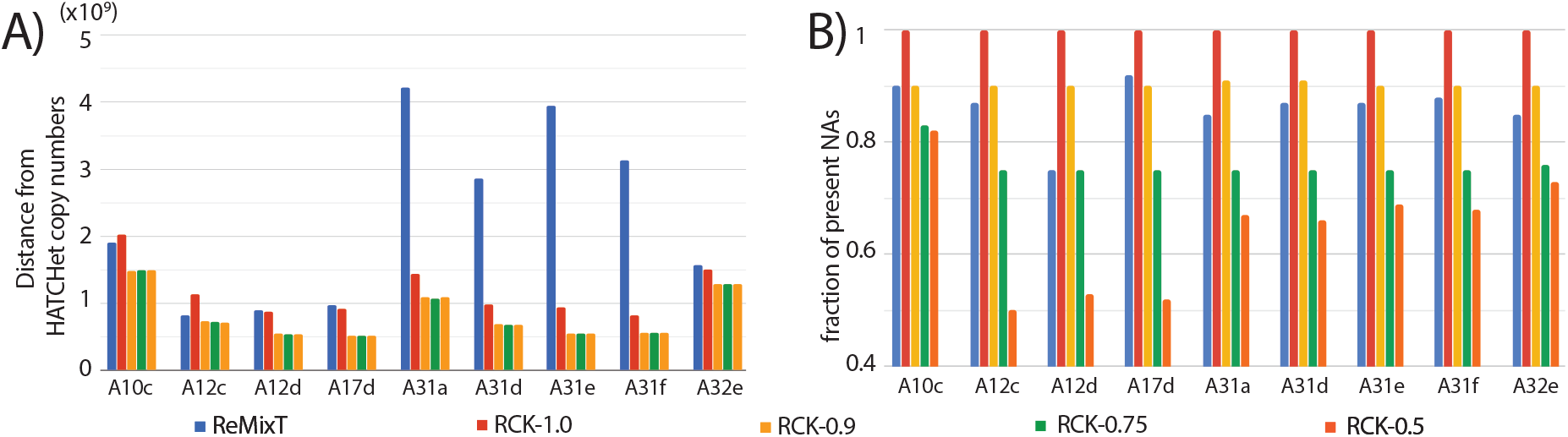
**A)** Length-weighted segment copy number distances (eq. (15)) between segment copy numbers from `HATCHet`and segment copy numbers output by `ReMixT`and `RCK`. **B)** Fractions of novel adjacencies (NAs) from input that are inferred to be present by `ReMixT`or `RCK`for each sample in the heterogeneous group. `RCK`used segment copy numbers from `HATCHet`in input and novel adjacency utilization parameter `P`= 1.0, 0.9, 0.75, 0.5.

We next compared the fraction of input novel adjacencies that were contained in the genomes reconstructed by `ReMixT`and `RCK`. This value ranged from 0.75 to 0.92 for `ReMixT`(Figure 2B). In contrast, for `RCK`fraction of utilized input novel adjacencies ranged from 0.5 to 1.0 with its lower bound explicitly controlled via the `P`parameter. We observe that `RCK`frequently utilizes more novel adjacencies than the minimum required (value of `P`). This occurs on 6/9 cancer samples (`A10c`, `A31a`, `A31d`, `A31e`, `A31f`, `A32e`) with `HATCHet`copy numbers and `P`= 0.75, `P`= 0.5, and 5/9 samples with `Battenberg`input. `RCK`’s incorporation of novel adjacencies at a higher proportion than the minimum required fraction `P`suggests that `RCK`is selectively including those novel adjacencies required to achieve copy number balance.

Next, we analyzed the structure of karyotypes inferred by each method. Since `ReMixT`does not output `A`/`B`labels for extremities involved in novel adjacencies, we investigated whether it was possible to derive `A`/`B`labels on `ReMixT`adjacencies to produce reasonable cancer genomes that would allow for a copy number balance/excess on the extremities of segments and comply with generalized IS constraints. We first observed that the karyotypes reconstructed by `ReMixT`had a large number of extremities that are not telomeres in the reference and have copy number excess (ranging from 41 to 133 per genome), corresponding to a large number of novel telomeres (Figure S1). Such karyotypes correspond to unlikely cancer genomes having dozens or even hundreds of linear chromosomes with novel telomeres, in addition to ∼46 (∼92 in WGD samples) derived linear chromosomes with reference telomeres. In contrast, the `RCK`results reported here use only reference telomeres and thus the karyotypes have at most 48 (89 in WGD samples) linear chromosomes in total.

We examined the frequency of violations of the generalized IS constraints. By construction, `RCK`karyotypes have no such violations. In contrast, we identified three types of violations of generalized IS conditions in the `ReMixT`karyotypes. The first is an intra-genome violation of the **homologous-extremity-exclusivity** constraint. This violation occurs when the inferred segment copy numbers require that a novel adjacency *a* be assigned *both* a label `A`and a label `B`in order to achieve copy number balance (Figure 3A). This situation requires that at least two large-scale somatic rearrangements occur independently at the same genomic position on both homologous chromosomes, which is highly unlikely. We find that karyotypes reconstructed by `ReMixT`contain such violations in 6/9 samples, ranging from 1 to 8 violations per genome, and from 1 to 12 violations per sample (Figure 3B).

**Figure 3:**
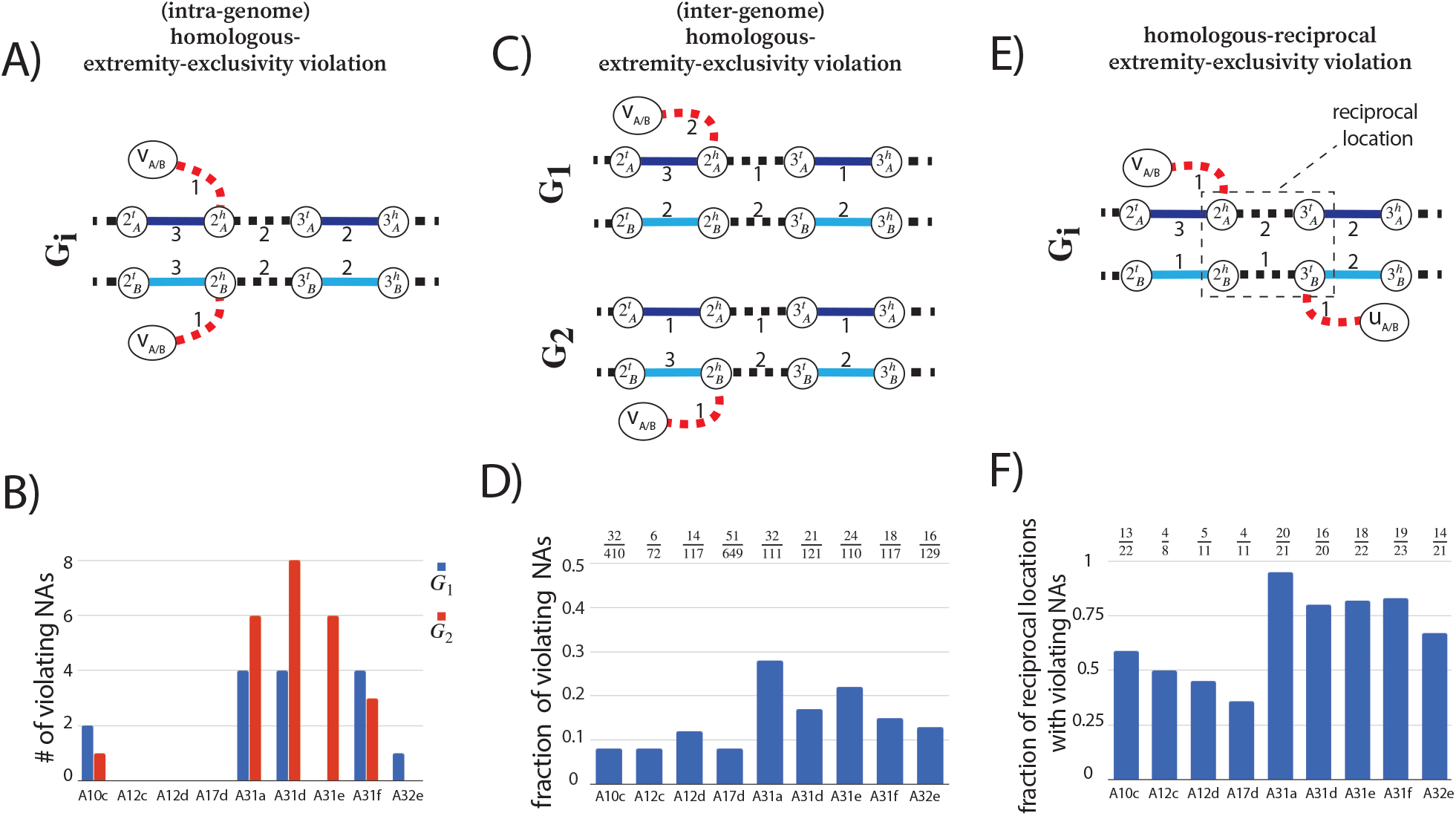
`ReMixT` karyotypes from the heterogeneous group of metastatic prostate cancer samples have numerous violations of the generalized IS constraints. In graph panels A), C), and E) solid edges represent segment edges, black-dashed edges represent reference adjacency edges, and red-dashed edges represent novel adjacency edges. Integer values indicate copy numbers of corresponding segment and adjacency edges. **A)** Example of a violation of the **homologous-extremity-exclusivity** constraint. To obtain copy number balance, both homologous vertices 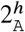 and 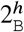 must be involved in novel adjacencies. **B)** Number of novel adjacencies (NAs) in each cancer karyotype inferred by `ReMixT` in each sample that violate the **homologous-extremity-exclusivity** constraint. **C)** Example of a violation of the inter-genome **homologous-extremity-exclusivity** constraint. To obtain copy number balance, both homologous vertices 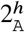 and 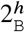 (in different genomes) must be involved in novel adjacencies. **D)** Fractions ^x^/_y_ of the number (*x*) of novel adjacencies (NAs) violating the inter-genome **homologous-extremity-exclusivity** constraint (on at least one of the extremities involved in every novel adjacency) in `ReMixT` karyotypes, over the total number (*y*) of novel adjacencies reported by `ReMixT` as being present in both genomes in the every sample. **E)** Example of a violation of the intra-genome **homologous-reciprocal-extremity-exclusivity** constraint. To obtain copy number balance, both homologous-reciprocal vertices 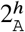 and 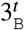 must be involved in novel adjacencies. We note that in addition to the violation of the intra-genome **homologous-reciprocal-extremity-exclusivity**) constraint, violations of the inter-genome or both version(s) of this constraint are also possible (See Figure S7). **F)** Fractions ^x^/_y_ of the number (*x*) of reciprocal locations with violations of either intra- or inter-genome (or both) **homologous-reciprocal-extremity-exclusivity** constraint in `ReMixT` karyotypes, over the total number (*y*) of reciprocal locations with both involved NAs reported as being present by `ReMixT`.

The second type of violation is an inter-genome violation of the **homologous-extremity-exclusivity** constraint. This violation occurs when a novel adjacency *a* is reported as being present in more than one genome in the sample, but a label `A` must be assigned to at least one *a*’s extremities in in one genome, and a label `B` must be assigned to at least one *a*’s extremities in another genome. This situation requires at least two large-scale somatic rearrangements occur *independently* at the same homologous genomic location in two different tumor clones, which is highly unlikely. We found that the karyotypes produced by `ReMixT` had such violations in all samples, with a substantial fraction (ranging from 0.09 to 0.28) of novel adjacencies containing such violations (Figure 3D).

The third type of violation concerns pairs of reciprocal novel adjacencies. For a pair *a* = {*x, uh*}, *b* = {(*u* + 1)*t, y*} of reciprocal novel adjacencies that involve reference adjacent extremities *uh,* (*u*+1)*t* possible violations of generalized IS include intra/inter-genome violation of the **homologous-extremity-exclusivity** or intra/inter-genome violation of the **homologous-reciprocal-extremity-exclusivity** constraints, or both. Any such violation requires that at least two large-scale somatic rearrangements occur *independently* on the same or homologous genomic location both producing pairs of reciprocal novel adjacencies, a situation which is highly unlikely. We found that karyotypes produced by `ReMixT` had such violations in all samples; furthermore in 6/9 samples more than half of reciprocal novel adjacencies had such violations (Figure 3F).

#### 2.2.3 Homogeneous tumor samples

We ran `RCK` and `ReMixT` on cancer samples from the homogeneous group and analyzed the karyotypes output by both methods, following the procedures described above for the heterogeneous samples. Since `ReMixT` assumes that an input sample contains exactly two cancer clones, `ReMixT`’s results disagree with both `Battenberg`’s and `HATCHet`’s predictions of one cancer clone in these samples. To obtain a partial comparison of the segment copy number profiles inferred by `ReMixT` with the profiles inferred by `Battenberg` and `HATCHet` in each sample, we used `ReMixT`’s clone with the highest cellular frequency. Overall, our analysis of inferred cancer genomes karyotypes in the homogeneous group aligned with the findings for the heterogeneous group. In particular, we found that on every sample in the homogeneous group, the segment copy numbers inferred by `RCK` (with `P` ≤ 0 9) are more similar to the copy numbers from `Battenberg` (Figure S6A) and `HATCHet` (Figure S6B) compared to the segment copy numbers inferred by `ReMixT`. We also found that the fraction of input novel adjacency that were present in inferred karyotypes ranged from 0.82 to 0.94 in `ReMixT` results and from 0.5 to 1.0 in `RCK` results (Figure S5). As in the case of heterogeneous samples, we observed that segment copy number distances are largest for `RCK` when we require `RCK` to use all novel adjacencies (a larger proportion than used in `ReMixT`), but the distances decrease and stabilize when some novel adjacencies are excluded (`P` ≤ 0 9).

Similar to the heterogeneous samples, we observed that karyotypes inferred by `ReMixT` had implausible features including a large number (and multiplicity) of novel telomeres (Figure S3) and violations of the generalized infinite sites constraints (Figures S4). In contrast, karyotypes inferred by `RCK` had no such issues. Overall, our analysis of inferred cancer genomes karyotypes in the homogeneous group aligned with the findings for the heterogeneous group.

## 3 Discussion

We presented `RCK`, a novel algorithm for reconstructing clone- and haplotype-specific cancer genomes karyotypes from bulk tumor samples. `RCK` accounts for heterogeneity in the observed tumor sample, correctly models the diploid reference genome, and enforces biologically reasonable evolutionary constraints that generalize the infinite sites constraints to somatic large-scale rearrangements. `RCK` is, to the best of our knowledge, the only algorithm with these features and also the only algorithm that can combine both next- and 3rd-generation sequencing data into the reconstruction process, leveraging the long-range adjacency information from 3rd-generation sequencing technologies.

On real cancer sequencing data, we found that `RCK` infers cancer karyotypes which inferred segment copy numbers are closer to those produced by state-of-the-art copy number inference tools (`HATCHet` and `Battenberg`), and which novel adjacencies conform with constraints from an infinite sites evolutionary model. In contrast, `ReMixT`’s approach of using novel adjacencies to “adjust” copy numbers generally led to allele-specific segment copy numbers that were different from those of `HATCHet` and `Battenberg`. Moreover, the novel adjacencies that are present in `ReMixT` inferred karyotypes often require biologically implausible rearrangements. These results demonstrate that “linking” of copy numbers via novel adjacencies without considering the underlying somatic evolutionary process is not advisable.

While the proposed `RCK` method uses a very comprehensive somatic evolutionary model and addresses several shortcomings of the previous approaches, there are limitations and avenues for future improvements. First, in the `RCK` results presented here, we assume that no new telomeres are introduced in the cancer genomes, i.e. all telomeres are telomeres of the reference genome. `RCK` allows for non-reference telomeres to be specified; however, we have not incorporated telomere selection into the objective function of the optimization. Such novel telomeres can correspond to real telomeres, but in many cases are likely due to missing novel adjacencies in the input data. Second, we can further generalize `RCK` to simultaneously analyze multiple samples from the same individual, perhaps including a phylogenetic [62] or longitudinal constraints [38]. Simultaneously analysis of multiple samples has proved useful in copy number inference [63]. Third, it would be helpful to model a patient-specific germline genome that includes germline structural variations, long repetitive segments, etc. Finally, one could further leverage information in 3rd-generation sequencing data by including haplotype-specific labeling of extremities involved in groups of novel adjacencies.

`RCK`’s inference of clone- and haplotype-specific cancer karyotypes enables further studies of the somatic mutational processes that produce highly rearranged cancer genomes, as well as improved characterization of specific functional changes (e.g., loss of heterozygosity, novel haplotype-specific fusion genes, etc). Higher-resolution reconstructions of cancer karyotypes can also help researchers illuminate differences/similarities between different types of cancer in general and lead to a more targeted and personalized medical treatments in specific patients.

## 4 Methods

We start by considering the case of “perfect” input data and the problem of reconstructing karyotype of a single mutated genome in sections 4.1 – 4.2. Then we extend to the case of a heterogeneous cancer sample (section 4.3). and describe how our model can incorporate (when available) information from 3rd-generation sequencing technologies (section 4.4). In section 4.5 we describe a more general case where there is uncertainty in input segment copy number values.

### 4.1 Single derived genome

We view cancer as a process propagated by a sequential application of somatic large-scale rearrangements starting with a *diploid* reference genome R and ending with a derived genome G. Every chromosome in a diploid reference genome R is present in two homologous copies, which we label by `A` and `B` respectively. A *segment* 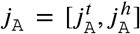 is a contiguous part of a reference chromosome labeled `A`; its endpoints 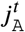 and 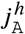 are called *extremities*. We label segments 1 through *m* in a multichromosomal diploid reference genome R. In a mutated genome that is derived from the reference via large-scale rearrangements, segments can be absent, present more than once, and appear both in forward and reverse orientation. We denote by 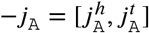 a reversed instance of segment *j*_A_.

Extremities that demarcate the beginning and the end of a chromosome are called *telomeres* and we define by τ (G) the set of telomeres in genome G.

A pair (*j*_A_, *k*_B_) of consecutive segments on a chromosome determines an *adjacency* 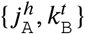 (i.e., a pair of extremities that are adjacent on a chromosome). A genome G determines a set(G) of adjacencies present in it.

For a diploid reference genome R with *k* chromosomes we define a set 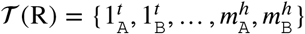 of reference telomeres and note that|𝒯 (R) |= 4*k*. We further note that a multichromosomal diploid reference R determines a set𝒜 (R) of *reference adjacencies* as follows:

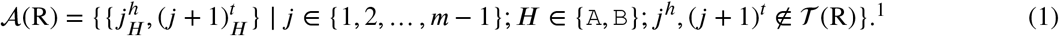

A derived genome G corresponds to a collection of concatenation of segments (i.e., derived chromosomes), where segments in each novel concatenation can originate from any homologous copy of any of the chromosomes in the diploid reference R. Each derived chromosome thus corresponds to a word from the following alphabet:

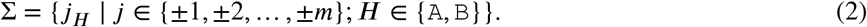

Adjacencies that are present in a mutated genome G but are not present in the reference are called *novel* and we denote by 𝒜*N* (G) a set of novel adjacencies in genome G. We note that since there are no novel adjacencies in the reference we have 𝒜 *N* (R) = Ø. We say that a set 𝒜 of adjacencies satisfies infinite sites if no two adjacencies in 𝒜 involve the same extremity. For a reference adjacency 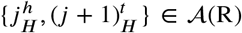 we call extremities 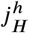 and 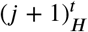 *reciprocal*.

We assume that the large-scale somatic rearrangements that “break” and “reglue” chromosomes do not affect the same genomic locations (on either of the or `A`/`B` copies) more than once during the entire somatic evolutionary process (i.e., generalized infinite sites). We note that under generalized IS only reference adjacencies can participate in breaks, however we also note that novel adjacencies produced by rearrangements can further be amplified/deleted via other rearrangements. For a break *r* of an adjacency 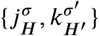 involving extremities 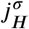 and 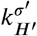 under the generalized IS at every point before and after *r* in the somatic evolutionary process none of the reference/novel adjacencies involving either 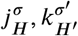 or 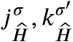 can be involved in any other rearrangement(s) (breaks), where *H, H ′ ∈* {`A`, `B`}, Â = `B`, and 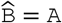. Examples of rearrangements that violate the generalized IS and consecutive implications for novel adjacencies in the derived genomes are shown in supplementary Figure S9.

With the generalized IS assumption for somatic evolution propagated by large-scale rearrangements we naturally obtain several constraints for the derived genome which we list below:

a. **extremity-exclusivity**: every extremity 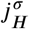 is involved in *at most* one novel adjacency from 𝒜_*N*_ (G). This constraint is based on the fact that for a novel adjacency *a* to involve an extremity 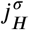 there must have been a large-scale rearrangement breaking a reference adjacency involving 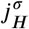 in the first place (and possible several other reference adjacencies). Having more than 1 novel adjacency involving 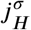 would correspond to the scenario where some other rearrangement must have broken some adjacency involving 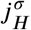, which is prohibited under generalized IS.
b. **homologous-extremity-exclusivity**: if an extremity 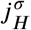 is involved in a novel adjacency from 𝒜 _*N*_ (G), then the homologous extremity 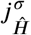 is *not* involved in any novel adjacency from 𝒜 _*N*_ (G). This constraint follows the logic outlined in **extremity-exclusivity**, but considers `A`/`B` labeled homologous extremities 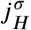and 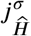 : for both 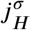 and 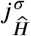 extremities to be involved in novel adjacencies there must have been at least two large-scale rearrangements breaking homologous reference adjacencies involving both extremities 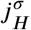 and 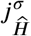, which is prohibited under the generalized IS.
c. **homologous-reciprocal-extremity-exclusivity**: if an extremity 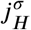 from the reference adjacency 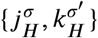 is involved in a novel adjacency from 𝒜 _*N*_ (G), then the homologous extremity 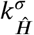, is *not* involved in any novel adjacency from 𝒜 _*N*_ (G). This constraint follows the justification provided in **homologous-extremity-exclusivity** for both extremities 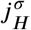 and 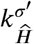 to be involved in novel adjacencies there must have been two large-scale rearrangements breaking both homologous reference adjacencies 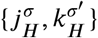 and 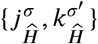 which is prohibited under the generalized IS.

We call a genome G *proper*, if the above three conditions are met.

A genome G determines a diploid segment copy number profile **C**_G_ = (**a** = [*a*_1_, *a*_2_, …, *a*_*m*_], **b** = [*b*_1_, *b*_2_, …, *b*_*m*_]), where values (*a*_*j*_, *b*_*j*_) ∈ ℕ^2^ indicate the number of copies of segments *j*_A_ and *j*_B_ in G. We note that in a diploid reference R we have *a*_*j*_ = *b*_*j*_ =1 for every segment *j*. An example of a diploid segment copy number profiles **C**_G_ and **C**_R_ for a derived genome G and a reference R are shown in Figure 4.

**Figure 4:**
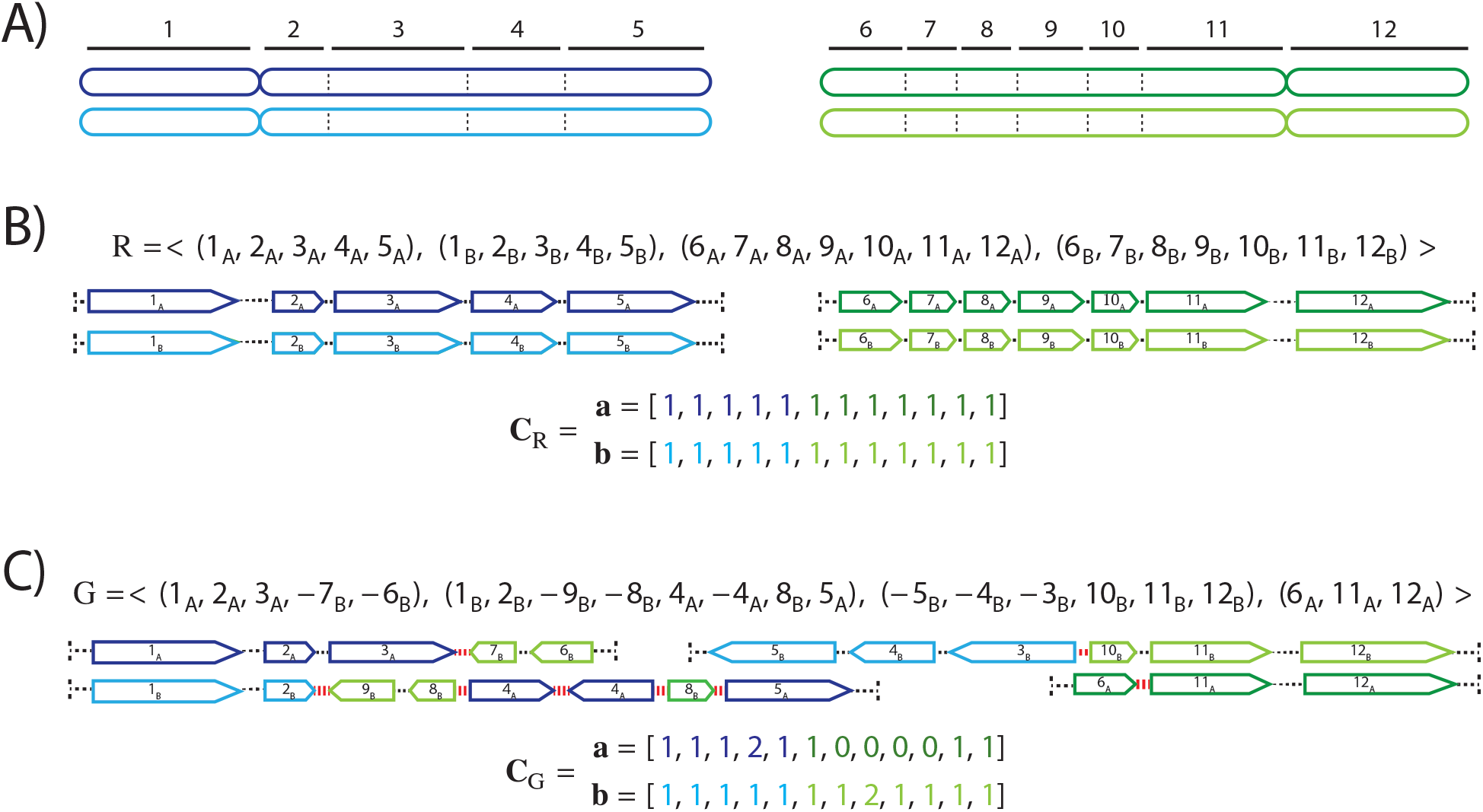
**A)** Example of a diploid reference genome R containing two pairs of homologous chromosomes (chromosomes labeled by `A` are shown in dark blue/green, and homologous copies labeled by `B` are shown in light blue/green) that are “partitioned” into 12 consecutive segments labeled 1 through 12. **B)** Reference genome R is shown as a collection of concatenations of segments, with segments located on chromosomes labeled `A` shown in dark blue/green and segments located on chromosomes labeled `B` shown in light blue/green. The “pointy” end of each segment *j* correspond to extremity *jh*, while the “flat” end corresponds to extremity *jt*. Dashed lines determine adjacencies between segments’ extremities. A set 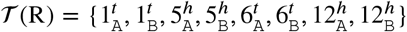 corresponds to telomeres in the shown diploid reference genome R. A diploid segment copy number profile **C**_R_ = (**a**, **b**) is shown for the genome R with colors (dark/light blue/green) corresponding to `A`/`B` labeled segments. **C)** A derived genome G obtained via multiple large-scale rearrangements from the reference genome R. Red dashed lines correspond to novel adjacencies (e.g., 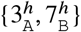). A diploid segment copy number profile **C**_G_ = (**a**, **b**) is shown for the genome G with colors (dark/light blue/green) corresponding to `A`/`B` labeled segments. A set 𝒯 (G) = 𝒯 (R) of telomeres in the derived genome G equals to that in the original reference genome R.

When a mutated genome G is derived from a diploid reference R current technologies do not allow us to measure its diploid segment copy number profile **C**_G_ directly. Rather there exist several methods [57, 9, 7, 40, 24, 19, 35, 63] that are capable of measuring a pair 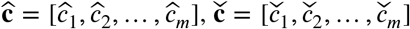,of vectors, where for every segment *j* an unlabeled (allele-specific) pair 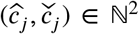 represents copy numbers of segments *j*_A_ and *j*_B_ in G, but without `A`/`B` labels explicitly associated with the measured values. In other words, we know that 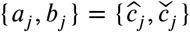, but it is unclear whether 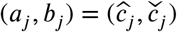 or 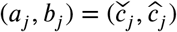 (example shown in Figure 6).

**Figure 5:**
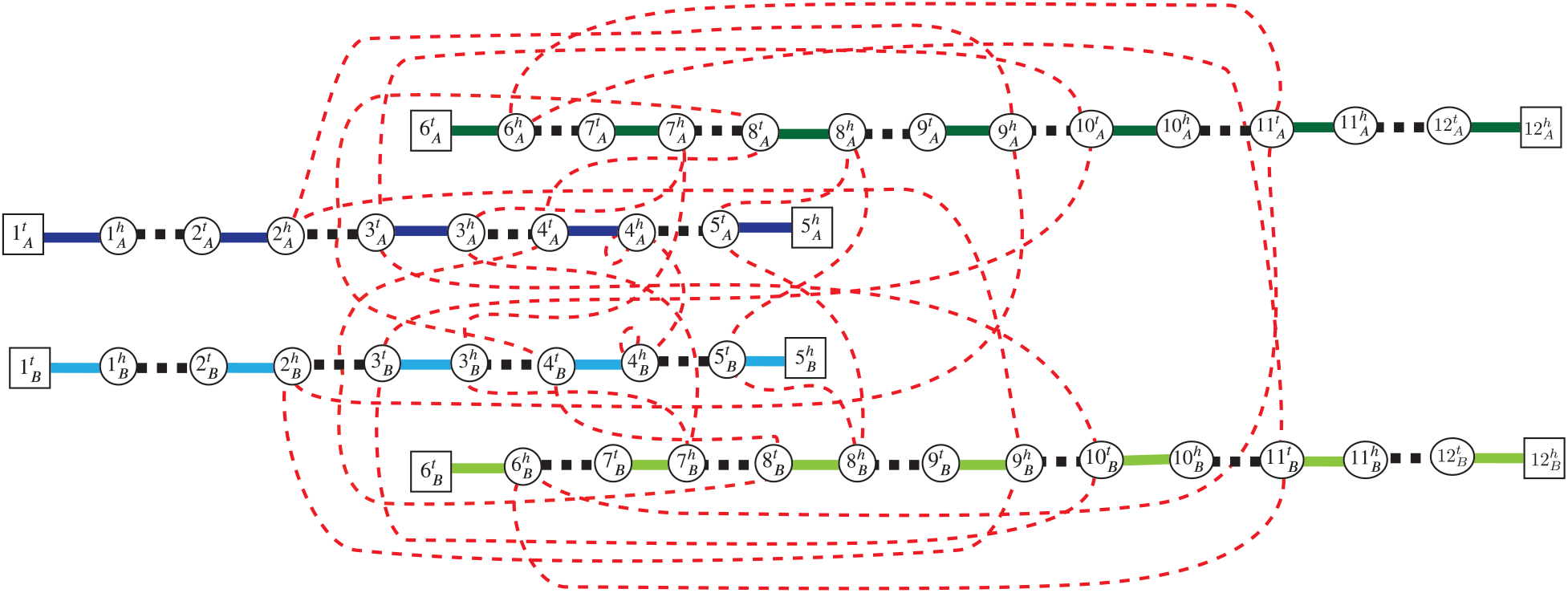
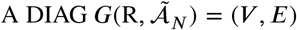 constructed on a set {1, 2, …, 12} of segments, and a set 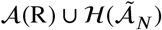 of adjacencies, where a set 𝒜(R) corresponds to reference adjacencies in a diploid reference R shown in Figure 4B, and a set 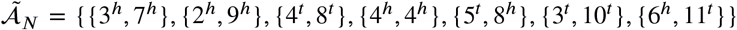 represents unlabeled novel adjacencies that were measured from a derived genome G shown in Figure 4C. Telomere vertices 𝒯 (*G*) =𝒯(R)⊆*V* are shown as squares, and non-telomere vertices are shown as circles. Solid edges correspond to segment edges in *E*_*S*_, with dark blue/green edges corresponding to segments labeled `A`, and light blue/green edges corresponding to segments labeled `B`. Reference adjacency edges *E*_*R*_ are shown as black-dashed edges, and novel adjacency edges *E*_*N*_ are shown as red-dotted edges.

**Figure 6:**
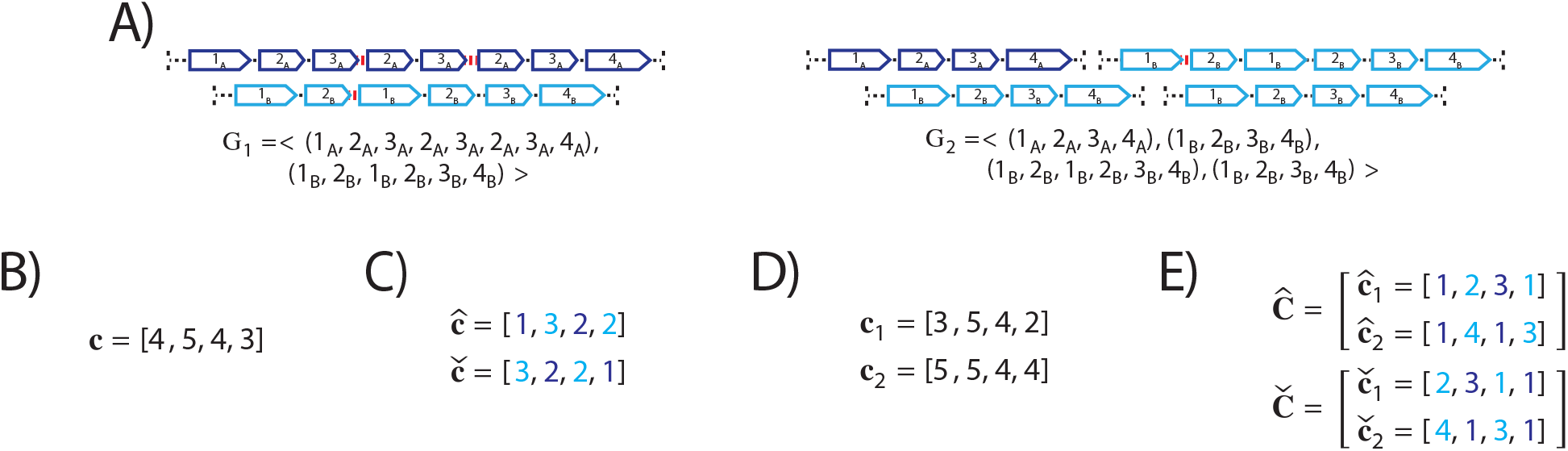
Description of errors in noise-free segment copy number (SCN) inference for a heterogeneous (i.e., 2 genomes) and haplotype-specific (i.e., A/B labeled segments) sample *S* = (G_1_, G_2_) under different limiting assumption about the sample’s structure. **A)** A 2-genome proper sample *S* = (G_1_, G_2_) with every genome G_*i*_ *∈ S* depicted both as collections of adjacent blocks as well as corresponding sequences of signed block. **B)** SCN inference under the assumption that the sample in question is homogeneous (i.e., comprised of a single derived genome) and with no consideration given to the fact that every segment has two distinct `A`/`B` instances of it (*haploid-reference*). In a vector **c** = [*c*_1_, *c*_2_, *c*_3_, *c*_4_] for a segment *j* a value *c*_*j*_ corresponds to an average over sums 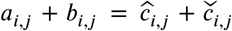 of diploid/allele-specific SCNs across genomes G_*i*_ *∈ S*. **C)** SCN inference under the assumption that the sample is homogeneous, but distinguishing between `A`/`B` labeled copies of every segment, though not preserving the alleles labels mapping to true `A`/`B` labels across segments. Colors encode true labeling (dark blue – `A`, light blue – `B`), *flipped* alleles are shown for segments 2 and 4). In vectors 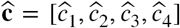 and 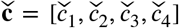 for a segment *j* values 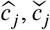 correspond to averages 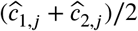 and 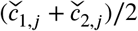 of genome- and allele-specific copy number values. **D)** SCN inference under the assumption that the sample is heterogeneous, but with a haploid-reference assumption. In vectors **c**_1_ = [*c*_1,1_, *c*_1,2_, *c*_1,3_, *c*_1,4_] and **c**_2_ = [*c*_2,1_, *c*_2,2_, *c*_2,3_, *c*_2,4_] for a segment *j* and genome G_*i*_ the value *c*_*i,j*_ equals to the sum 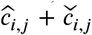 of allele-specific copy number values in a genome G_*i*_. **E)** Allele- and genome specific SCN inference. Colors encode true labeling (dark blue – `A`, light blue – `B`), flipped alleles are shown for segment 2 and 4 (i.e., 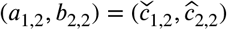 and 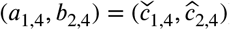).

Furthermore, when a genome G derives from a diploid reference R we can not measure a set *A* (G) of adjacencies in G directly, but rather we can only measure an obfuscated version of the set 𝒜 _*N*_ (G). of novel adjacencies in G. That is, for every novel adjacency 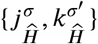 (G) we can only measure an *unlabeled*(i.e., with involved extremities missing the `A`/`B` labels) adjacency 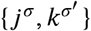 (e.g., for a derived genome G shown in Figure 4 instead of measuring a novel adjacency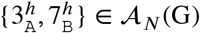. we measure an unlabeled novel adjacency {3^*h*^, 7^*h*^}). There exist several methods capable of producing the unlabeled novel adjacencies both from a standard short-read bulk sequencing data [51, 48, 30, 10, 60] as well as from 3rd-generation sequencing technologies [49, 16, 52, 66, 50, 27, 17].

We note that if a set 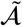 of unlabeled novel adjacencies is measured from a proper derived genome, it satisfies the generalized infinite sites conditions: since in unlabeled novel adjacencies involved extremities lack `A`/`B` labels, only the (unlabeled) **extremity-exclusivity** constraint (i.e., on unlabeled extremities) must be satisfied, which is achieved, because in proper genome conditions **extremity-exclusivity** and **homologous-extremity-exclusivity** guarantee that for every pair 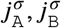 of homologous extremities at most one of them is involved in any novel adjacency from *𝒜*_*N*_ (G), and thus the unlabeled extremity *j* is also involved in at most one measured unlabeled novel adjacency from 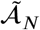.

We assume that large-scale rearrangements that generated a mutated genome G from a diploid reference R have not created novel telomeres (i.e., 𝒯 (G) ⊆ 𝒯 (R)), and formulate the following problem of reconstructing mutated genome from measurement data:

**Problem 1.** *Given a diploid reference* R, *allele-specific copy number measurements* 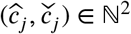 *for every segment j, and a set* 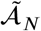 *of unlabeled novel adjacencies that satisfies (unlabeled)* ***extremity-exclusivity*** *constraint, find a proper genome* G *satisfying:*

1. *for every adjacency* 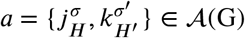 *either* 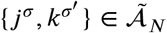 *or a* ∈ *A*(R);
2. *for every adjacency* 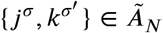 *there exist labels H, H′* ∈ {*`A`, `B`*}, *such that* 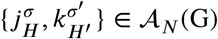;
3. *for every* (*a*_*j*_, *b*_*j*_) ∈ **C**_G_ *either* 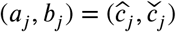 or 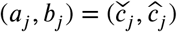
4. *𝒯 (G) ⊆ 𝒯 (R).*

Since the measured unlabeled novel adjacencies do not have the `A`/`B` labels, we do not know the true underlying novel adjacencies that produced a measurement. For an unlabeled novel adjacency 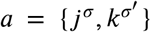 we defined by 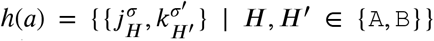 a set of the four possible novel adjacencies that can be obtained by `A`/`B` labeling extremities in *a*. For a given set 𝒜 of unlabeled novel adjacencies we define a set ℋ (𝒜) of all possible novel adjacencies as follows:

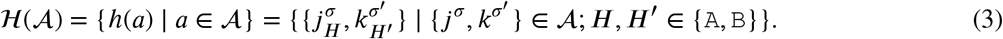

We note that when a set 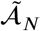 of measured unlabeled novel adjacencies comes from a genome G, it follows that 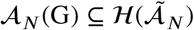. A union 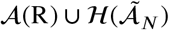 of sets 𝒜 (R) and 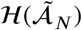 represents all possible adjacencies that can be present in the observed mutated genome G.

### 4.2 Diploid Interval Adjacency Graph

We reformulate Problem 1 of finding a proper derived genome G from the measurement data as a graph-theoretic problem. First, we define the *diploid interval adjacency graph* (DIAG), which can be viewed as a generalization of a breakpoint graph used in the area of comparative genomics [1, 65, 3], or graphs used in the area of structural analysis of normal and cancer genomes with haploid reference structure [36, 42, 32, 12, 15, 35. 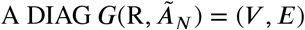 is constructed on a set {1, 2, …, *m*} of segments, and a set 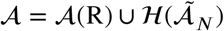 of adjacencies.

The set *V* of vertices is in one-to-one correspondence with all segments’ extremities. Formally we define *V* as follows:

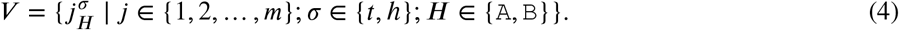

The set *E* of edges in a DIAG is comprised of two types of edges: *segment* edges *E*_*S*_ and *adjacency* edges *E* _*𝒜*_. The set *E*_*S*_ of segment edges represents segments as follows:

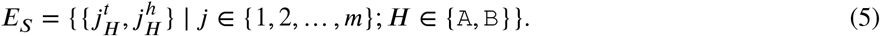

The set *E*_*𝒜*_ of adjacency edges is in a one-to-one correspondence with a set 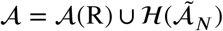 of adjacencies: i.e., every adjacency 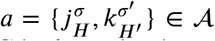 is represented by a corresponding adjacency edge 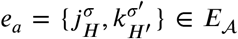 (e.g., an example DIAG is shown in Figure 5).

Every adjacency edge *e*_*a*_ ∈ *E* _*𝒜*_ that corresponds a reference adjacency *a* ∈𝒜 (R) we call a *reference adjacency edge*, and we denote by *E*_*R*_ ⊆ *E* _*𝒜*_ a set of all reference adjacency edges in *E* _*𝒜*_. We also define a set *E*_*N*_ = *E* _*𝒜*_ \ *E*_*R*_ of *novel adjacency edges*, with edges in *E*_*N*_ respectively corresponding to novel adjacencies in 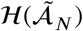. Since adjacency edges and adjacencies are in one-to-one correspondence we allow ourselves to use adjacencies when referring to adjacency edges and vice versa.

Since every vertex 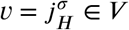 is incident to exactly one segment edge 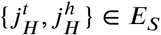, we define *e*_*S*_ (*v*) ∈ *E*_*S*_ to be a segment edge incident to a vertex *v*, and define *e*_*s*_ (*j*_*H*_) ∈ *E*_*S*_ to be a segment edge corresponding to a segment *j*_*H*_. Every vertex *v* ∈ *V* is incident to at most one reference adjacency edge, and we define *e*_*R*_(*v*) ∈ *E*_*R*_ to be a reference adjacency edge containing vertex *v*, if such adjacency exists. Naturally, we define *E*_*N*_ (*v*) ⊆ *E*_*N*_ to be a set of novel adjacency edges incident to *v* ∈ *V*.

Every chromosome in a derived genome G determines a segment-adjacency edge alternating walk in the corresponding DIAG, that starts and ends at telomere vertices in 𝒯 (G) (examples are shown in the supplement Figure S8B). Such an alternating walk spells out a concatenation of segments from the reference genome, corresponding to a derived chromosome in G. Thus, a derived genome G determines a collection of segment-adjacency edge alternating walks. The number of times a segment edge 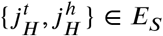 is traversed (in either direction) across all walks determined by G corresponds to the segment copy number (e.g., 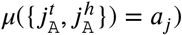). Similarly, the number of times an adjacency edge 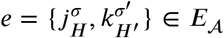 is traversed (in either direction) across all walks determined by G corresponds to an *adjacency copy number* (i.e., the number of times an adjacency corresponding to an edge *e* is present in G). A genome G thus determines an *edge multiplicity function μ* : *E* → 𝒩 on both segment and adjacency edges (example is shown in the supplement Figure S8A). We call the corresponding DIAG *G*(R, *𝒜*_*N*_, *μ*) a *weighted DIAG*.

We note that DIAG is allowed to have self-loop adjacency edges that correspond to a self-loop novel adjacencies in 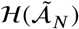. Such self-loop novel adjacencies can be produced by breakage-fusion-bridge cycles, inverted tandem duplications, and other more complex large-scale genome rearrangements that have been observed in cancer [22, 64, 33, 25]. We define by *l*(*a*) : *E*_𝒜_→{1, 2} an auxiliary function that outputs 2 if *a* is a self-loop adjacency (edge), and 1 otherwise. We say that a vertex *v* ∈*V* exhibits a *copy number balance* provided:

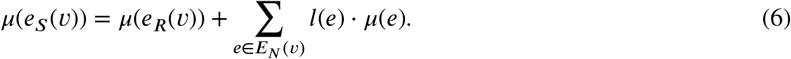

Similarly, we say that a vertex *v* ∈ *V* exhibits a *copy number excess* provided:

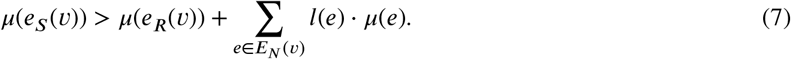

The following theorem follows directly from previous work [29, 45]:

**Theorem 1.** *A weighted DIAG G* = (*V, E, μ*), *can be decomposed into a collection of segment-adjacency edge alternating walks that start and end at a set* 𝒯⊆*V of telomere vertices, such that every edge e* ∈ *E is traversed μ*(*e*) *times, if:*

1. *every non-telomere vertex v* ∈ *V* \𝒯 *is copy number balanced,*
2. *and every telomere vertex v* ∈ 𝒯 ⊆*V has a copy number excess.*

When the derived genome is allowed to have circular chromosomes, which have been extensively observed and studied in cancer [8, 21, 59, 18, 56], Theorem 1 provides not only a necessary, but also a sufficient condition for a derived genome to exist. For an extended discussion about DIAG decomposition into segment-adjacency edge alternating walks please refer to supplementary material section S2.1.

For every unlabeled novel adjacency 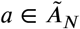 and a 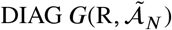 we define by *h*^*E*^ (*a*) ⊆*E*_*N*_ a subset of novel adjacency edges corresponding to adjacencies in *h*(*a*). Furthermore, given a weighted 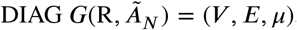, for every unlabeled novel adjacency 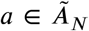 we define by 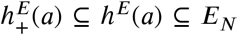 a subset of adjacency edges with positive multiplicities as follows:

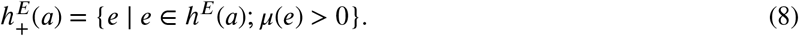

Now we readily reformulate the Problem 1, allowing a derived genome to contain circular chromosomes, into a problem of finding edge multiplicities in the associated DIAG as follows:

**Problem 2.** *Given a* 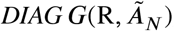, *where the set* 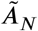 *of unlabeled novel adjacencies satisfies (unlabeled)* ***extremity-exclusivity*** *constraint, and allele-specific copy number measurements* 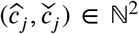 *for every segment j of telomere vertices find an edge multiplicity function μ* : *E → N such that:*

1. *for every unlabeled adjacency* 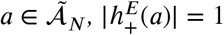;
2. *for every self-loop unlabeled adjacency* 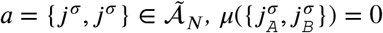;
3. *for every pair*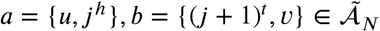 *of unlabeled novel adjacencies, such that* 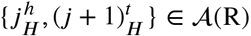, *there exist* 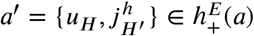 *and*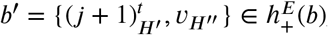, *where H,H ′,H ″* ∈ {*`A`, `B`*};
4. *for every segment j, either* 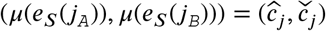 *or* 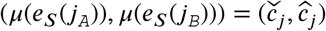;
5. *every non-telomere vertex v*∈*V* \𝒯(R) *exhibits copy number balance* (eq. (6));
6. *every telomere vertex v* ∈𝒯(R) ⊆*V exhibits either copy number balance* (eq. (6)*) or copy number excess* (eq. (7)).

We note, that finding an edge multiplicity function *μ* in Problem 2 guarantees the existence of a proper derived genome that determines *μ*, but such derived genome does not necessarily need to be unique. A resulting weighted DIAG *G* = (*V, E, μ*) thus determines a haplotype-specific karyotype of the derived genome in question.

### 4.3 Multiple derived genomes

The sequencing assays in cancer genomics can involve biological samples that can be genetically heterogeneous (i.e., comprised of cells with different derived genomes, also sometimes referred to in the literature as *clones*). Let us assume that a sample (i.e., set of genomes) *S* = (G_1_, G_2_, …, G_*n*_) in question is comprised of *n* genomes all of which have derived from a diploid reference R via large-scale rearrangements. A sample *S* = (G_1_, G_2_, …, G_*n*_) determines a pair **C**_*S*_ = (**A** = [**a**_1_, **a**_2_, …, **a**_*n*_]^*T*^, **B** = [**b**_1_, **b**_2_, …, **b**_*n*_]^*T*^) of *n ×m* diploid segment copy number matrices, where genome-specific segment copy number vectors **a**_*i*_ = [*a*_*i,*1_, *a*_*i,*2_, …, *a*_*i,m*_] and **b**_*i*_ = [*b*_*i,*1_, *b*_*i,*2_, …, *b*_*i,m*_] contain integer values *a*_*i,j*_, *b*_*i,j*_ ∈ℕ that correspond to the number of times segments *j*_A_ and *j*_B_ appear in genome G_*i*_ ∈ *S* respectively. We denote by **A**_[*j*]_ = [*a*_1,*j*_, *a*_2,*j*_, …, *a*_*n,j*_]^*T*^ and by **B**_[*j*]_ = [*b*_1,*j*_, *b*_2,*j*_, …, *b*_*n,j*_]^*T*^ vectors of copy number values for segments *j*_A_ and *j*_B_ across all genomes G_*i*_ ∈*S*.

For a sample *s* = (G_1_, G_2_, …, G_*n*_) we do not measure the pair **C**_*S*_ = (**A**, **B**) of its *n ×m* diploid segment copy matrices directly, but rather we measure a pair 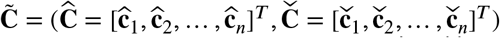 of *n ×m* allele-specific segment copy number matrices, such that for every segment *j* either 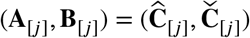 or 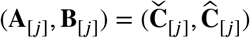. Examples of allele-specific vs diploid alongside other errors in a noise-free segment copy number inferences with different limiting assumptions about the sample’s structure are shown in Figure 6.

For a sample *S* = (G_1_, G_2_, …, G_*n*_) we define by 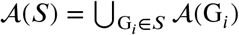 a set of all adjacencies and by 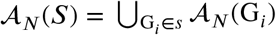 a set of all novel adjacencies present in any (subset) of the genomes in *S*.

Similarly to the case of a single derived genome, out ability to measure novel adjacencies from a sample *S* = (G_1_,G_2_, …, G_2_) is obfuscated. For every novel adjacency 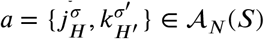 we can only measure an unlabeled counterpart 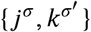 and we also loose the information about which genome(s) in sample *S* the underlying novel adjacency *a* is actually present in. We define by 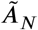 a set of unlabeled adjacencies measured from a sample *S*.

We generalize the previously introduced constraints on possible structures of the derived genomes G_*i*_ ∈*S* for the sample *S*. We call a sample *s* = (G_1_, G_2_, …, G_*n*_) *proper* if the **extremity-exclusivity**, **homologous-extremity-exclusivity**, and **homologous-reciprocal-extremity-exclusivity** assumptions hold, with a set 𝒜_N_ (G) substituted with a set 𝒜_N_ (*S*) (i.e., considering a set 𝒜_N_(*S*) of novel adjacencies across all of the genomes in the observed sample S). Substituting 𝒜_*N*_(G) with 𝒜_*N*_ (*S*) allows us to impose the generalized IS constraints for the whole somatic evolutionar process (i.e., take into account rearrangement that occur on all the branches of the somatic phylogenetic tree) that produced the observed sample S. We note that if a set 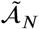 of unlabeled novel adjacencies comes from a roper sample *S*, then 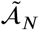 satisfies the generalized IS conditions (by satisfying the (unlabeled) **extremity-exclusivity** constraint). Moreover, we note that if a sample *S* = (G_1_, G_2_, …, G_*n*_) is proper, then any subsample (including individual derived genomes G_*i*_ *∈ S*) of *S* is also proper.

A generalized version of Problem 1 for a sample *S* = (G_1_, G_2_, …, G_*n*_) is stated below:

**Problem 3.** *Given a diploid reference* R, *a pair* 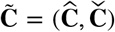 *of n* × *m allele-specific segment copy number matrices, and a set* 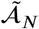 *of measured unlabeled novel adjacencies that satisfies (unlabeled)* ***extremity-exclusivity*** *constraint, find a proper sample s* = (G_1_, G_2_, …, G_*n*_) *such that:*

1. *for every adjacency* 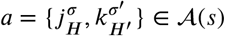, *either* 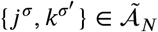 *or* 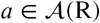
2. *for every adjacency* 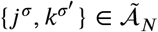, *there exists a unique pair* 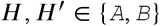 *of labels, such that* 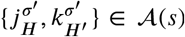;
3. *for every segment j, either* 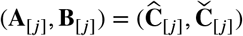 *or* 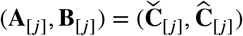;
4. *for every genome* G_*i*_ *∈ s, the telomere set* τ (G_*i*_) *⊆* τ (R).

In a sample *S* = (G_1_, G_2_, …, G_*n*_) and a set 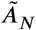 of unlabeled novel adjacencies measured form *S*, we observe a DIAG 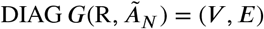. Every genome G_*i*_ ∈ *S* determines a genome-specific edge multiplicity function *μ*_*i*_ : *E* ⟶ ℕ as was previously described in a case of a single derived genome.

We extend previously introduced copy number balancing conditions (6) and (7) on vertices in *V,* using genome-specific edge multiplicity functions. For a genome G_*i*_ ∈ *S*, a vertex *v* ∈ *V* exhibits *copy number balance* provided:

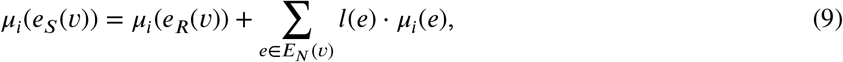

and a vertex *v* ∈ *V* exhibits *copy number excess* provided:

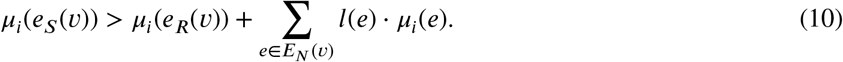

For every unlabeled adjacency 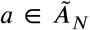 and a genome G_*i*_ ∈ *S* we define by 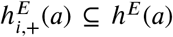 a subset of novel adjacency edges in *h*^*E*^ (*a*) with positive copy number as determined by the genome-specific edge multiplicity function *µ*_*i*_ as follows:

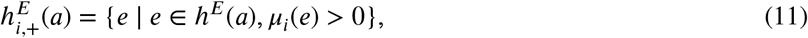

and we then naturally generalize the definition of 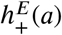 for the sample *S* = (G_1_, G_2_, …, G_*n*_) case:

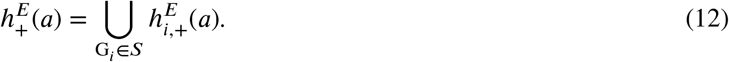

For every segment *j*_*H*_ we define by ***µ***_[*j,H*]_ = [*µ*_1_(*e*_*S*_ (*j*_*H*_)), *µ*_2_(*e*_*S*_ (*j*_*H*_)), …, *µ*_*n*_(*e*_*S*_ (*j*_*H*_))]^*T*^ a vector of genome-specific edge multiplicity functions’ values on the segment edge *e*_*S*_ (*j*_*H*_) ∈ *E*_*S*_.

We now reformulate a general Problem 3 of finding a proper sample *S* = (G_1_, G_2_, …, G_*n*_) in terms of finding edge multiplicity functions *µ*_1_, *µ*_2_, …, *µ*_*n*_ : *E* ⟶ ℕ in the corresponding DIAG as follows:

**Problem 4.** *Given a DIAG G*(R, *Ã*_*N*_) = (*V, E*), *where a set Ã*_*N*_ *of unlabeled novel adjacencies satisfies (unlabeled)* ***extremity-exclusivity*** *constraint, and a pair* 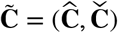 *of n* × *m allele-specific segment copy number matrices, find edge multiplicity functions µ*_1_, *µ*_2_, …, *µ*_*n*_ : *E ⟶* ℕ *such that:*

1. *for every adjacency* 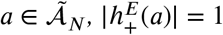;
2. *for every i* ∈ [*n*] *and every adjacency* 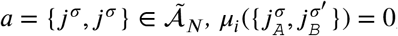;
3. *for every pair a* = {*u, j*^*h*^}, *b* = {(*j* + 1)^*t*^, *v*} *∈ Ã*_*N*_ *of unlabeled novel adjacencies, such that* 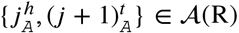, *there exists* 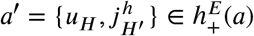 *and* 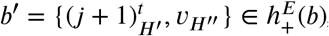, *where* ***H, H*′**, ***H*″** ∈ {*A, B*};
4. *for every segment j, either* 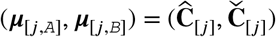 *or* 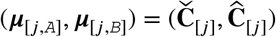;
5. *for every i* ∈ [*n*] *and every non-telomere vertex v* ∈ *V* \ 𝒯 (R) *the equality* (9) *holds;*
6. *for every i* ∈ [*n*] *and every telomere vertex v* ∈ 𝒯 (R) ⊆ *V either the equality* (9) *or the inequality* (10) *hold.*

### 4.4 3rd generation sequencing technologies and novel adjacency groups

Besides the cost-efficient next-generation sequencing technologies (i.e., bulk-sequencing with short paired-end reads), there exist other, more expensive, 3rd-generation sequencing technologies (e.g., single-cell, barcoded linked reads, and long-read sequencing) that can provide additional insight about measured unlabeled novel adjacencies [16, 52, 66, 49, 50, 27, 17]. We observe a sample *S* = (G_1_, G_2_, …, G_*n*_) and a set *Ã*_*N*_ of unlabeled novel adjacencies coming from *S*. We define a *3rd-generation sequencing experiment* as either all reads obtained in a single-cell sequencing essay, a set of reads annotated with the same barcode in the barcoded sequencing experiment, or a single long read obtained by a long-read sequencing technology. Let us assume that a 3rd-generation sequencing experiment on a *S* = (G_1_, G_2_, …, G_*n*_) identifies a group *u* ⊆ *Ã*_*N*_ of unlabeled novel adjacencies. Since every 3rd-generation sequencing experiment is conducted either on a single cell (e.g., single-cell) and thus produces data from a single derived genome, or on a part of a single derived chromosome (e.g., bar-codded, long-range) present in a single derived genome, the group *u* of unlabeled adjacencies is guaranteed to originate from a single derived genome G_*i*_ ∈ *S*.

We note that for every unlabeled novel adjacency 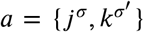 measured from a sample *S* = (G_1_, G_2_, …, G_n_) there exist a unique novel adjacency counterpart 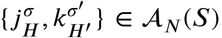, when *S* is proper. Or, more formally, |*h* (*a)* ∩𝒜_*N*_(*S*) |= 1. Thus, for every group *u*⊆ *Ã*_*N*_ of unlabeled novel adjacencies measured via a single 3rd-generation sequencing experiment on a proper sample *S* = (G_1_, G_2_, …, G_*n*_) for at least one genome G_*i*_ ∈ *S* we have:

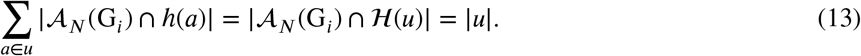

### 4.5 Uncertainty in copy number measurements

As we have stated before, there exist several methods that for a given sample *S* = (G_1_, G_2_, …, G_*n*_) aim to infer a pair 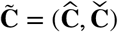 of *n* × *m* allele-specific segment copy number matrices. Loss of explicit information about `A`/`B` labels across segments in such inference (while preserving segment’s specific allele separation across all genomes in *S*) is often not the only limitation of these methods. It is also often the case for sequences of reference-adjacent segments to be grouped together into larger non overlapping *fragments*, for which the allele-specific copy numbers are inferred.

More formally, we call a sequence (*j,j*+1,….,*j*+*l*) of reference adjacent segments a *fragment* and denote it by *f*_[*j,l*]_. We denote by ℱ a collection of non overlapping fragments that cover all of the segments.

When allele-specific copy numbers are inferred on fragments, rather than individual segments, we naturally obtain the same copy number values for all segments within every overarching fragment, which may be incorrect for some or even all segments within the observed fragment. On the other hand, allele-specific nature of the inferred fragments copy numbers preserves allele separation not only across genomes, but also across segments within each fragment. We thus view available allele-specific copy numbers for fragments as an approximation of the true underlying segment copy numbers, and try to infer the true underlying diploid segment copy number values, while leveraging the allele separation preservation across segments within each fragment.

Let us observe a pair **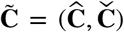** of *n* × *m* allele-specific segment copy number matrices, a pair **C** = (**A**, **B**) of *n* × *m* diploid segment copy number matrices, and a set ℱ of fragments. For every fragment *f* ∈ ℱ we define a length-weighted copy number distance **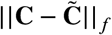** as follows:

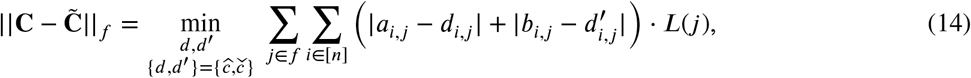

where *L*(*j*) is the total number of base pairs (i.e., length) of segment *j*. We further define a copy number distance **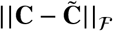** between pairs **C** and 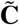 of diploid and allele-specific segment copy number matrices as follows:

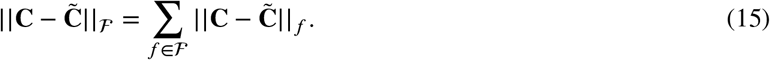

We now extend the previous Problem 3 of finding a sample from the measured data to the case when the measured allele-specific segment copy numbers are noisy and (optionally) information from 3rd-generation sequencing experiments is available:

**Problem 5.** *Given a diploid reference* R, *a pair* **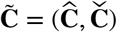** *of n* × *m allele-specific segment copy number matrices, a set ℱ of fragments, a set Ã*_*N*_ *of measured unlabeled novel adjacencies that satisfies (unlabeled)* ***extremity-exclusivity*** *constraint, and (optionally) a set 𝒰 of groups of unlabeled novel adjacencies, find a proper sample s* = (G_1_, G_2_, …, G_*n*_) *such that:*

1. *for every adjacency* 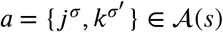 *either* 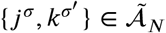 or *a* ∈ 𝒜(R);
2. *for every adjacency* 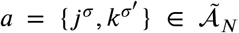 *there exists a unique pair H, H*′ ∈ {*`A`, `B`*} *of labels, such that* 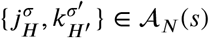;
3. *for every adjacency group u* ∈𝒰, *there exists (at least one) genome* G_*i*_ ∈ *s such that |𝒜*_*N*_ (G_*i*_) ∩ℋ (*u*) =|*u|;*
4. *for every genome* G_*i*_ ∈*s the 𝒯* (G_*i*_)⊆ *𝒯* (R); *and the copy number distance* 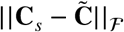 *is minimized.*

A reformulation of Problem 5 in terms of finding edge multiplicity functions on the edges of the corresponding DIAG is provided below:

**Problem 6.** *Given a* 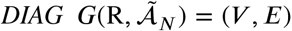, *where a set* 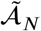 *of unlabeled measured novel adjacencies satisfies (unlabeled)* ***extremity-exclusivity*** *constraint, a (optionally) set* 𝒰 *of groups of unlabeled novel adjacencies, a pair* 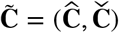 *of n* × *m allele-specific segment copy number matrices, and a set* ℱ *of fragments, find edge multiplicities functions μ*_1_, *μ*_2_, …, *μ*_*n*_ : *E* → ℕ *such that:*

1. *for every adjacency* 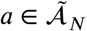 *we have* 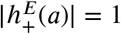;
2. *for every i* ∈ [*n*] *and every adjacency* 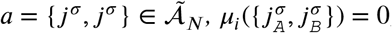;
3. *for every adjacency group u* ∈ 𝒰 *there exists (at least one) i* ∈ [*n*] *such that* 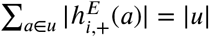;
4. *for every pair* 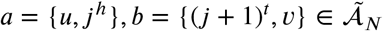 *of unlabeled novel adjacencies, such that* 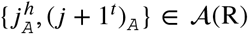, *there exists* 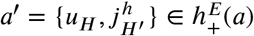 *and* 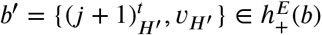, *where H, H′, H″* ∈ {*`A`, `B`*};
5. *for every i* ∈ [*n*] *and every non-telomere vertex v* ∈ *V* \ 𝒯 (R) *the equality* (9) *holds;*
6. *for every i* ∈ [*n*] *and every telomere vertex v* ∈ 𝒯 (R) ⊆ *V either the equality* (9) *or the inequality* (10) *hold; and such that for a pair* **C**_*μ*_ = (**A**_*μ*_, **B**_*μ*_) *of diploid segment copy number matrices (determined by values of edge multiplicity functions μ*_1_, *μ*_2_, …, *μ*_*n*_ *on segments edges E*_*S*_*), the copy number distance* 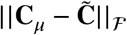 *is minimized.*

In the Supplement, we derive a mixed integer linear program (MILP) optimization problem that solves Problem 6.

### 4.6 Deriving extremities and novel adjacencies from data

Segment copy number inference methods often define a fixed-size partition of the reference genome into segments and thus constrain the coordinates of segments extremities. Every measured unlabeled novel adjacency determines a pair {(`chr`_1_, `coord`_1_, `str`_1_), (`chr`_2_, `coord`_2_, `str`_2_)}, where `chr`_*i*_determines the chromosome of origin the genomic loci *i*, `coord`_*i*_ determined the coordinate of the genomic loci *i* on the respective chromosome `chr`_*i*_, and `str`_*i*_ ∈ {+, -} determined the strand of origin of the genomic loci *i*.

Extremities of segments that are inferred by methods that measure clone- and allele-specific segment copy numbers and those involved in measured unlabeled novel adjacencies do not always align. Moreover, there is often a small uncertainty in the exact values of the coordinate `coord`_*i*_ of the genomic loci *i* involved in a novel adjacencies.

We first address the issue of refining the positions of extremities involved in reciprocal novel adjacencies. For every sample *S* we first observe all unlabeled novel adjacencies *Ã*_*N*_ measured from *S* and sort the positions involved in adjacencies from *Ã*_*N*_ on every chromosome (in descending order of the `coord`values). Then, using a sliding window approach, we update the coordinates for any consecutive pair *p*_*i*_, *p*_*j*_ of positions which resembles a reciprocal signature: i.e., if the distance | `coord`_*i*_ - `coord`_*j*_ | was less than 50 base pairs and `str`_*i*_ ≠ `str`_*j*_, we update the values of the coordinates in positions *p*_*i*_ and *p*_*j*_ so that they have a coordinate distance of 1, with the position having a + strand appearing prior to the position having a - strand (Figure 7A).

**Figure 7:**
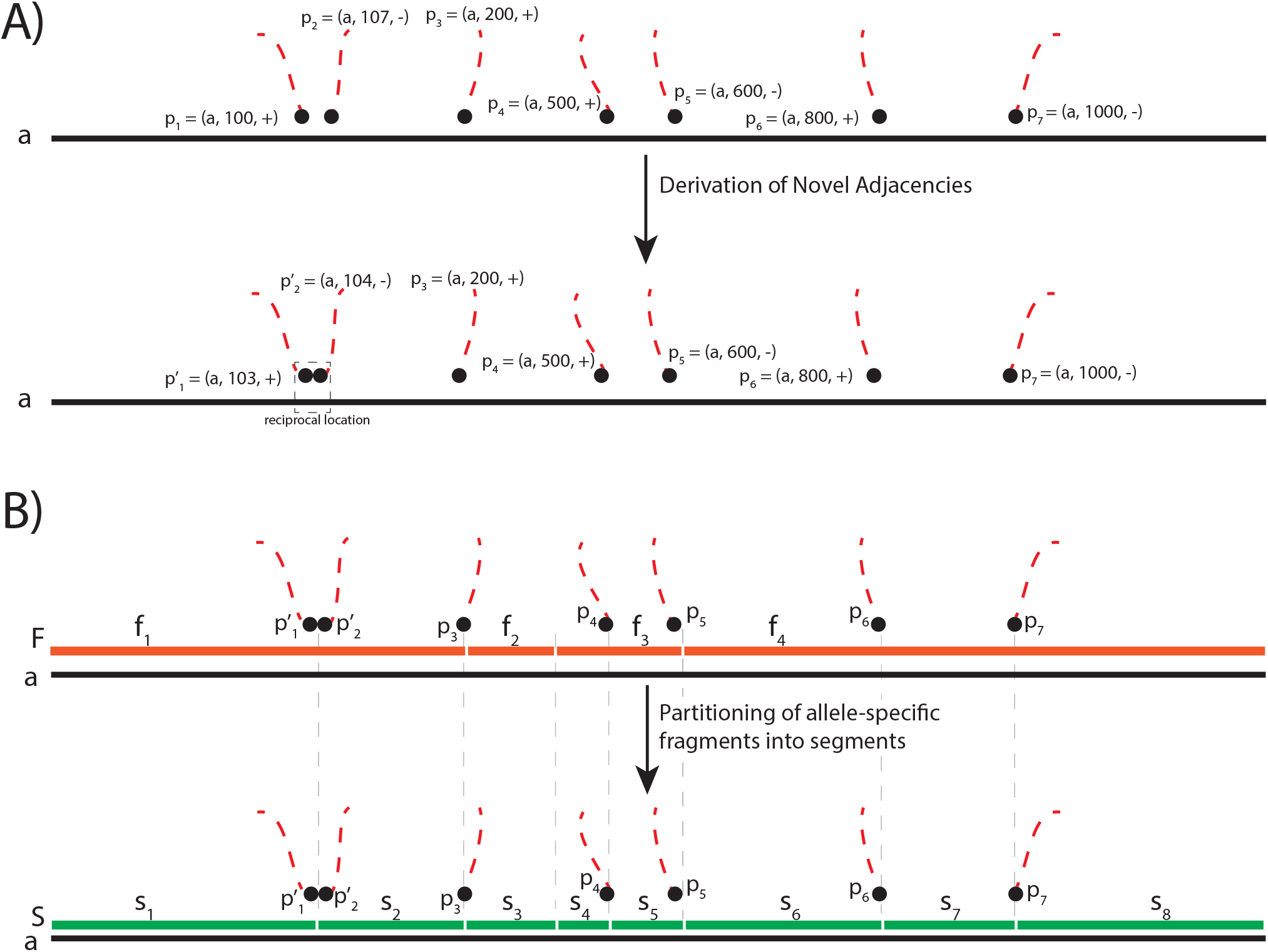
Derivation of the input for ReMixT and RCK. **A)** An example of derivation of coordinates that resemble a reciprocal signature in measured unlabeled novel adjacencies on a chromosome *a*. Positions *p*_1_ = (*a,* 100, +) and *p*_2_ = (*a,* 107, -) have reciprocal signature (i.e., | `coord`_1_ - `coord`_2_ | = 7 < 50 and `str`_1_ = – ≠ `str`_2_ = +). Updated pair 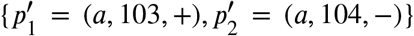 of coordinates constitutes a reciprocal location. **B)** An example of partitioning of a set ℱ = {*f*_1_, *f*_2_, *f*_3_, *f*_4_} of fragments from allele-specific copy number calls into a set ***S*** = {*s*_1_, *s*_2_, *s*_3_, *s*_4_, *s*_5_, *s*_6_, *s*_7_, *s*_8_} of segments. Extremities of segments in ***S*** correspond to either preprocessed coordinates of unlabeled novel adjacencies (e.g., 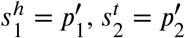) or to the extremities of fragments in ℱ (e.g. 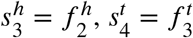).

Then, for allele-specific segment copy number input (e.g., from `Battenberg`and `HATCHet`) we partition fragments, on which allele-specific copy number values are measured, into smaller segments such that extremities of obtained segments either correspond to the coordinates of extremities involved in the preprocessed novel adjacencies from *Ã*_*N*_, or to the extremities of the original fragments (Figure 7B). Copy numbers on newly obtained segments are inherited from the values of the “parent” fragments.

Lastly, in order to compute length-weighted segment copy number distances between `RCK`, `ReMixT`, `Battenberg`, and `HATCHet`inferences on the prostate cancer samples, we refined the fragments/segments on which the copy numbers were inferred as demonstrate in Figure S10).

## Acknowledgments

This work is supported by a US National Institutes of Health (NIH) grants R01HG007069 and U24CA211000 and US National Science Foundation (NSF) CAREER Award (CCF-1053753) to BJR.

## Code availability

`RCK` is available on GitHub at https://github.com/raphael-group/RCK.

## Footnotes

1 We assume that every segment appears exactly once in a forward orientation the reference genome.

